# Multi-scale reactor designs extend the physical limits of CO_2_ fixation

**DOI:** 10.1101/2024.08.28.610213

**Authors:** Amir Akbari, Bernhard O. Palsson

## Abstract

CO_2_ valorization is a promising strategy for climate adaptation and transitioning towards a circular carbon economy. Here, we present a multi-scale, integrated systems approach for designing biomanufacturing systems that can utilize CO_2_ as a feedstock, focusing on the Wood–Ljungdahl and reductive glycine pathways. This approach relies on first principles, coupling the optimization of pathway and process variables. We examine the CO_2_-fixation capacity of both pathways in single- and multi-compartment reactor systems, demonstrating that the reductive glycine pathway has the potential to fix CO_2_ at significantly higher rates than photosynthetic organisms. We show that small differences in the energy-dissipative and stoichiometric structures of carbon-fixation pathways could significantly impact optimal designs and feasible design spaces. Our first-principle, systems-level approach quantifies these differences and uncovers strategies to expand the design space and extend the physical limits of carbon fixation, offering insights into pathway selection and process configurations for efficient biomanufacturing.

## Introduction

Mitigating the rising levels of atmospheric CO_2_ is a pressing global challenge. CO_2_ is the predominant greenhouse gas emitted through human activity and is recognized as the largest contributor to global warming [1]. A circular carbon economy is a promising manufacturing paradigm that can address this challenge by minimizing reliance on fossil fuels and reducing waste [2]. However, efficient conversion of CO_2_ into desired products remains a significant challenge due to its inherent stability and low energy content [3]. Overcoming these hurdles requires interdisciplinary approaches integrating techniques from metabolic engineering, synthetic biology, and process systems engineering.

Carbon-neutral manufacturing through biological means is a common strategy for climate adaptation [4]. C_1_ carbon sources (*e.g*., CO_2_, formate, methanol, and methane) can be converted to valuable goods through natural or synthetic biochemical pathways [5]. Leveraging the biosynthetic capabilities of natural autotrophs is an obvious choice for carbon fixation [6]. In this regard, chemoautotrophs are especially of interest because their carbon and energy metabolism have evolved to directly reduce CO_2_ using simple reductants, such as hydrogen. They are also more energy-efficient than photoautotrophs in general [5, 7, 8]. However, these non-model microorganisms have special nutrient demands and offer limited genetic tools, rendering them undesirable for industrial biotechnology [6, 9], although there has recently been progress with acetogens in large-scale production of industrial chemicals [10].

An alternative approach is to implement C_1_-fixation pathways in model heterotrophs for which advanced genome editing tools are available [11]. The reductive glycine pathway [12–14], Calvin-Benson-Bassham cycle [15], and ribulose monophosphate cycle [16] are examples of natural C_1_-fixation pathways engineered in *E. coli* and *S. cerevisiae*. Synthetic pathways, such as the GED [17], THETA [18], and serine threonine cycles [19] have also been introduced in model organisms. However, these non-native pathways are thermodynamically unfavorable and prone to side reactions, which can cause redox and energy imbalances, diminishing the growth rate of the host [20].

To avoid the foregoing undesirable interactions and reduce interference from competing metabolic pathways, C_1_-fixation pathways can be implemented in a cell-free setting [21], notable examples of which are the CETCH [22], ASAP [23], and HOPAC [24] cycles. Cell-free systems are less constrained than cell factories, offer precise control over enzyme concentrations, and are flexible in enzyme selection [25]. However, cofactor regeneration remains a challenge for these systems [25]. Moreover, the CO_2_ assimilation rates achieved so far were within the same range as those of photosynthesis, and design optimization was limited to reaction and enzyme engineering [5].

Here, we develop a constraint-based approach for integrated design of carbon fixation pathways and biomanufacuring processes, providing a framework for systems-level optimization of C_1_ carbon systems (Fig. 1). This approach couples the optimization of variables at the pathway and process level, providing the best strategies to alleviate the fundamental bottlenecks of carbon fixation that are optimal in the context of a biomanufacuring process. We apply this framework to examine the capacity of the Wood–Ljungdahl and reductive glycine pathways—two of the most natural, energy efficient carbon fixation pathways [7]—to fix CO_2_ in single- and multi-compartment reactor systems, showing that the reductive glycine pathway can furnish several orders-of-magnitude higher assimilation rates than photosynthetic organisms. Analysis of single-compartment and multi-compartment process designs for the Wood–Ljungdahl and reductive glycine pathways quantitatively revealed how various strategies for pathway and process design optimization can expand the design space, extending the physical limits of carbon fixation.

**Figure 1:**
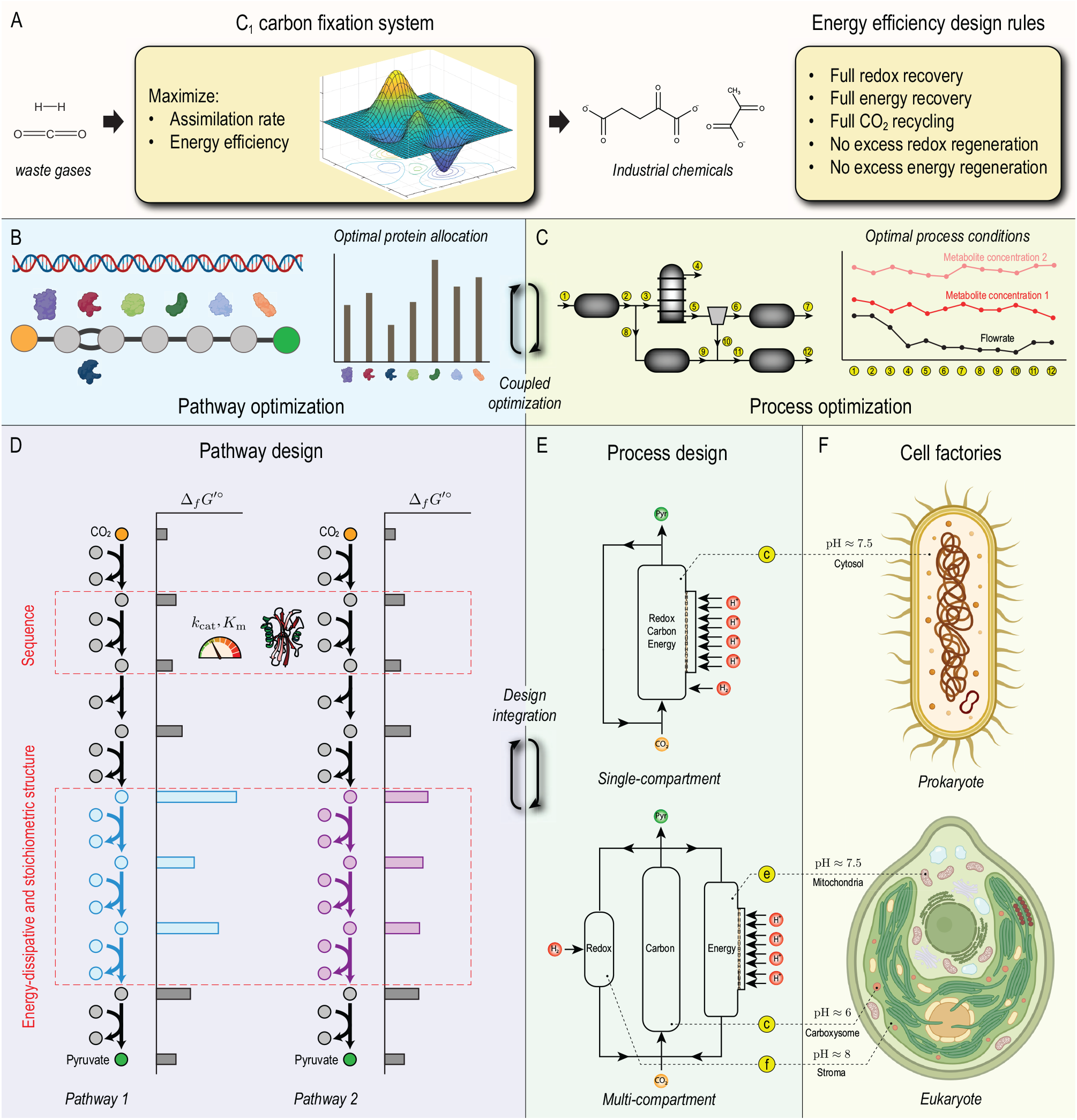
Computer-aided design of C_1_-fixation systems. (A) An optimization framework is developed to integrate pathway and process designs for economic production of industrial chemicals from C_1_ carbon sources. The objective is to maximize the CO_2_ assimilation rate and energy efficiency. Maximum energy efficiency is achieved indirectly by implementing a set of design rules within the formulation of the optimization problem (see “Design objectives”). This framework couples the optimization of carbon fixation pathways, furnishing their optimal functional states (*e.g*., protein allocation) and of (C) process systems, providing their optimal operating conditions (*e.g*., stream flow rates). It also enables the evaluation of (D) pathway design strategies (*e.g*., modulating stoichiometric and dissipative structures) and (E) process configurations (*e.g*., compartmentalization) in tandem. (F) Single- and multi-compartment reactor systems examined in this study are analogous to prokaryotic and eukaryotic cell factories.

## Results

### Multi-scale integrated systems approach

Carbon fixation is subject to biophysical constraints arising from bottlenecks in reaction energies and rates, substrate solubilities, and cofactor regeneration. These constraints are fundamental and limit the carbon-fixation capacity of both natural and synthetic pathways in cell factories or cell-free systems. Their underlying bottlenecks often lead to multi-scale trade-offs in C_1_-based manufacturing systems. Optimizing manufacturing processes with bottlenecks that interact across multiple time and length scales requires an integrated systems approach.

We introduce a multi-scale, systems-level framework to design C_1_-based manufacturing processes. We adopt a constraint-based approach that hinges on first principles to identify optimal designs, so it can be applied to biological and non-biological manufacturing systems in a unified way; this study, however, only focuses on biological, cell-free systems. We impose fundamental biophysical constraints, including (i) mass balance, (ii) kinetics, (iii) thermodynamics, (iv) ion binding, (v) Maxwell’s first law, and (vi) macro-molecular crowding. The goal is to design processes, and specifically reactor systems, in which C_1_ fixation occurs through enzymatic reactions. Optimal designs are determined by solving an optimization problem, where the objective is to maximize the C_1_ assimilation rate and energy efficiency (Fig. 1A). To simplify this multi-objective optimization problem, the energy-efficiency objective is achieved indirectly by imposing auxiliary constraints associated with a series of energy-efficiency rules (see “Design objectives” for details).

The C_1_-based design framework in this study integrates pathway and process design spaces, coupling the optimization of decision variables at the pathway (Fig. 1B) and process (Fig. 1C) level. It searches for manufacturing solutions in the integrated design space to simultaneously maximize pathway and process performance, while accounting for bottlenecks that interact at the pathway and process level. The integrated design space can generally be high-dimensional and is built from two subspaces associated with pathway and process variables. Variables in the ‘pathway subspace’ may be associated with protein sequences, pathway topology, pathway stoichiometric and dissipative structures, and enzyme concentrations (Fig. 1D), whereas those in the ‘process subspace’ may describe process operating conditions, process topology, equipment sizes, and unit-operation configurations (Fig. 1E). The design space in this framework is larger than those explored in previous studies on carbon fixation, where design optimization was based on pathway-search algorithms [17, 22], enzyme engineering [22], or reaction engineering [18]; thus, it provides more design alternatives to search through to enhance the assimilation rate and energy efficiency.

Although the framework outlined above is general, coupled optimization of all process and pathway variables is computationally challenging. Therefore, we prescribe the topology and structure of the pathway and process to simplify the analysis, only optimizing variables corresponding to the operating conditions of the process and functional state of the pathway.

At the pathway level, we fix the topology by choosing two natural carbon-fixation pathways rather than searching through the space of enzymatic reactions. Specifically, we examine the carbon-fixation capacity of the (i) Wood–Ljungdahl and (ii) reductive glycine pathways in a cell-free setting (Fig. 2). Both pathways are linear with CO_2_ the carbon source and pyruvate the product. They have the same topology but slightly differ in their stoichiometric and dissipative structures. To drive carbon fixation, these pathways require ATP and reducing cofactors, which are costly to replenish [25]. Therefore, to achieve a closed system that only relies on simple energy sources (proton gradient and hydrogen), we include ATP synthase and hydrogenases to regenerate ATP and reducing cofactors, respectively. At this level, metabolite concentration, enzyme concentrations, and metabolic fluxes are decision variables. However, turnover numbers and Michaelis constants are given parameters of optimization and are determined from kinetic constant databases (https://www.brenda-enzymes.org) (see “C_1_-CAD framework” for details). Although kinetic constants are not decision variables, their optimal values can be ascertained using parametric analysis and deep-learning based sequence optimization [26, 27].

**Figure 2:**
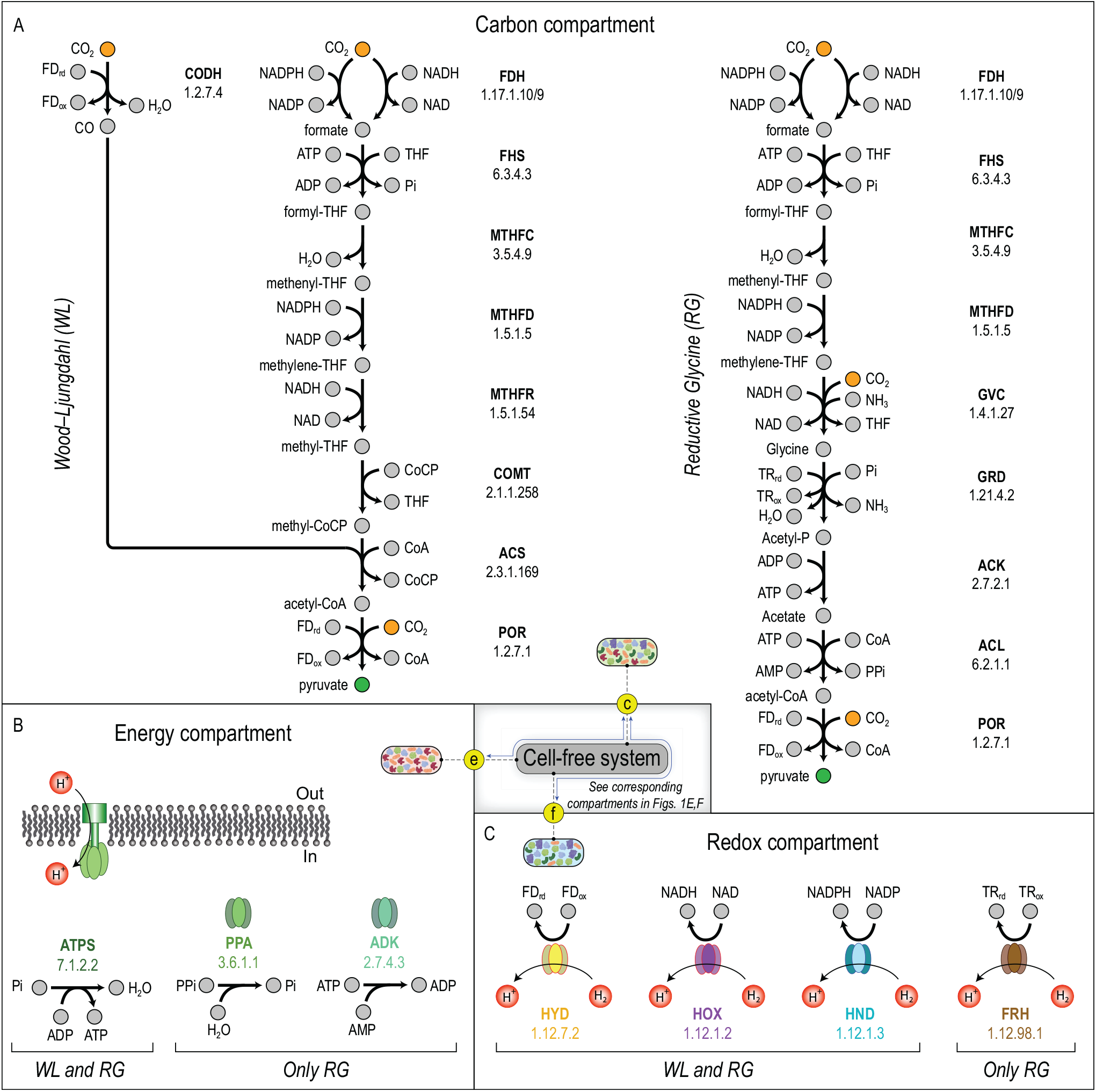
Design of a cell-free system for carbon fixation. (A) The CO_2_-fixation capacity of the Wood–Ljungdahl and reductive glycine pathways are examined for single- and multi-compartment reactor configurations. These pathways are among the most energy efficient natural carbon fixation pathways [7] and are, thus, promising candidates for the design of C_1_-fixation systems. Besides the main pathways involved in carbon metabolism, additional enzymes are required to regenerate the (B) energy-carrying and (C) reducing cofactors that drive CO_2_ fixation. In this work, hydrogenases and ATP synthase regenerate the reducing cofactors and ATP, respectively. In the single-compartment configuration, all enzymes involved in carbon metabolism and cofactor regeneration are in the same compartment, whereas in the multi-compartment configuration, enzymes involved in carbon metabolism, redox regeneration, and energy regeneration are separated and placed in specialized compartments termed carbon compartment, redox compartment, and energy compartment (see Figs. 1E,F).

At the process level, we fix the topology by considering two reactor configurations: (i) multi-compartment and (ii) single-compartment (Fig. 3A). In the first, enzymes are separated into a redox, carbon, and energy compartment according to the pH at which the associated reactions are thermodynamically favorable [28, 29]. Furthermore, a portion of the product stream from the carbon compartment is directed to the redox and energy compartment through recycle streams to replenish the redox and energy charge, respectively. In the second, all enzymatic reactions occur in the carbon compartment at the same pH. In both configurations, CO_2_ is pumped into the carbon compartment through the feed stream and pyruvate is extracted from the product stream. Moreover, all unreacted metabolites are recycled back into the carbon compartment. At this level, metabolite concentrations and flow rates in process streams are decision variables, while recycle ratios and pH are given parameters of optimization (see “C_1_-CAD framework” for details). The idea behind choosing the foregoing reactor configurations is to demonstrate how process design strategies can expand the space of feasible manufacturing systems, realizing solutions that otherwise would not be accessible.

**Figure 3:**
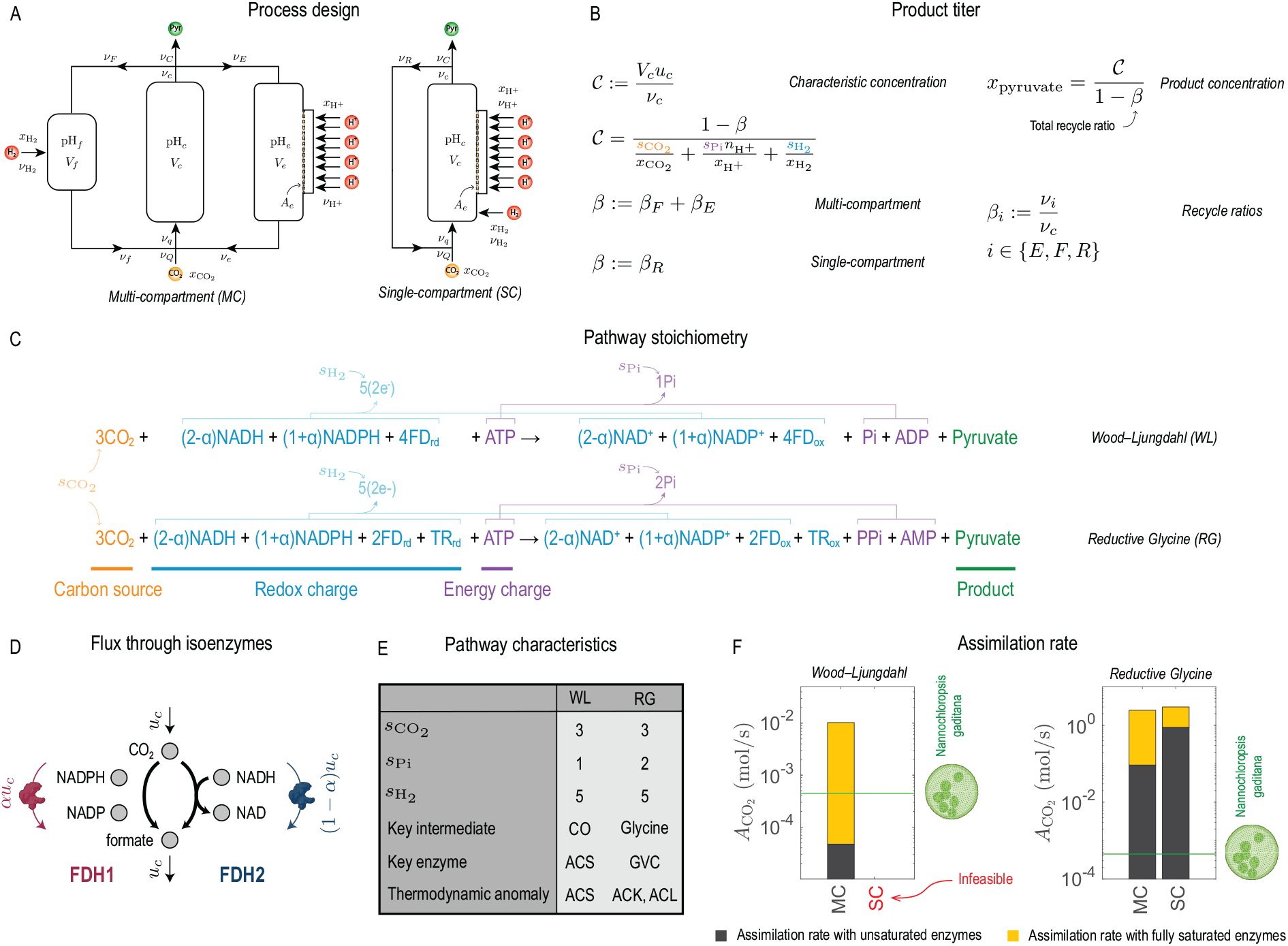
Coupled optimization of C_1_-fixation pathways and processes. (A) Process variables, process parameters, and boundary metabolite concentrations determining the optimal designs of single- and multi-compartment reactor configurations. Carbon-fixation and redox regeneration reactions respectively run in an acidic (pH_*c*_ *<* 7) and alkaline (pH_*f*_ *>* 7) environment in the multi-compartment configuration, while they all run at the same pH_*c*_ in the single-compartment configuration. (B) A characteristic concentration 𝒞 is defined in terms of *V*_*c*_, *ν*_*c*_, and *u*_*c*_, from which the product titer can be ascertained. Here, *V*_*i*_ denotes the volume of compartment *i, ν*_*i*_ the volumetric flow rate of stream *i, x*_*i*_ the concentration of boundary metabolite *i*, 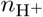 the number of protons translocated per ATP molecule in ATP synthase, and *u*_*c*_ the canonical flux through carbon metabolism. (C) The overall stoichiometry of the Wood–Ljungdahl and reductive glycine pathways and the definition of the stoichiometric coefficients *s*_*i*_ in (B). (D) Isoenzymes are handled by introducing a flux split ratio *α* for the corresponding reaction. (E) Comparison of pathway characteristics. (F) The maximum CO_2_-assimilation capacity of the Wood–Ljungdahl and reductive glycine pathways in the single-compartment (pH_*c*_ = 7.5) and multi-compartment (pH_*c*_ = 6) configurations. The upper bound *x*^ub^ = 0.05 M is imposed on metabolite concentrations in all process streams (see Table S1). Black bars indicate the assimilation rate if enzyme saturation efficiencies are accounted for, and yellow bars indicate how much the assimilation rate could be enhanced if all enzymes were fully saturated.

### Multi-scale optimization of carbon-fixation systems

We assessed the carbon-fixation capacity of the Wood–Ljungdahl and reductive glycine pathways in the multi- and single-compartment reactor configurations, evaluating the key metrics that determine the economy of the overall process: the rate, titer, and yield [30]. The energy-efficiency constraints imposed for design optimization ensure that no unreacted substrate leaves the reactor system. Therefore, the substrate is stoichiometrically converted to the product, so that the product yield is 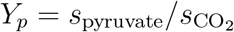 with *s*_pyruvate_ and 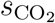 the stoichiometric coefficients of the product and substrate in the carbon-fixation pathway. These coefficients are identical for both pathways (Fig. 3C), thus the product yield is always *Y*_*p*_ = 1*/*3. Moreover, applying the energy-efficiency constraints results in a characteristic concentration 𝒞 that defines the product titer (Fig. 3B). This concentration depends on the stoichiometric coefficients, process parameters, and the concentration of boundary metabolites, so it is a constant and not affected by design optimization. Therefore, in this section, we only focus on the CO_2_ assimilation rate as a performance metric for alternative designs.

The structural characteristics of carbon-fixation pathways (Fig. 3E) largely determine the optimal design of the manufacturing system in which they are implemented. Stoichiometrically, the Wood–Ljungdahl and reductive glycine pathways yield the same number of pyruvate molecules for each fixed CO_2_ molecule, both requiring five NADH-equivalent of reducing cofactors and one ATP. However, the first cleaves one phosphate group from the ATP molecule spent and the second cleaves two. Energetically, these pathways are also different in their dissipative structures. A notable example is activated CO_2_ intermediates: besides formate, the key intermediate in the Wood–Ljungdahl pathway is CO, and in the reductive glycine pathway it is glycine. Because CO is a much higher energy intermediate than glycine, the Wood–Ljungdahl pathway has a more rugged formation-energy landscape than the reductive glycine pathway; thus, its feasibility is more restricted by thermodynamic constraints.

To alleviate the thermodynamic bottlenecks that restrict the feasibility of carbon-fixation pathways, we leveraged the pH-sensitivity of biochemical reactions [31, 32]. In general, reactions involved in carbon fixation, such as those of the Wood–Ljungdahl and reductive glycine pathways, are favorable in acidic environments, while redox regeneration reactions are favorable in alkaline environments. By compartmentalizing reactions that respond similarly to pH changes, we can extend the feasibility of carbon-fixation systems using design strategies at the process level. To test this idea, we evaluated the optimal CO_2_ assimilation rate of the two pathways in the multi- and single-compartment reactor configurations.

To determine how compartmentalization affects the maximum CO_2_-fixation capacity of carbon-fixation pathways, we optimized the design of both reactor configurations under the same conditions, except for pH. We imposed an upper bound on metabolite concentrations in all process streams within a physiological range (*x*^ub^ = 0.05 M). In the multi-compartment configuration, carbon-metabolism, redox-regeneration, and energy-regeneration enzymes were maintained at pH_*c*_ = 6, pH_*f*_ = 8, and pH_*e*_ = 7, respectively. However, in the single-compartment configuration, all the enzymes operated at pH_*c*_ = 7.5 (see Table S1). We then compared the CO_2_ assimilation rates with those of the photoautotroph *N. gaditana* [33]. Under these conditions, no steady state exists for the Wood–Ljungdahl pathway in the single-compartment configuration. However, in the multi-compartment configuration, a steady state can be achieved, but the CO_2_ assimilation rate is ∼10 times lower than in photosynthesis (Fig. 3F, black bar). We performed the same analysis assuming that all enzymes are fully saturated (see “Design optimization”) and found that the Wood–Ljungdahl pathway has the potential to fix CO_2_ at a significantly higher rate than photosynthesis if Michaelis constants are optimized (Fig. 3F, yellow bar). In contrast, the reductive glycine pathway is feasible in both reactor configurations and furnishes ∼10^2^–10^3^ times higher CO_2_ assimilation rates than in photosynthesis, depending on whether enzyme saturation efficiencies are accounted for. This high performance is due to the desirable dissipative structure of the reductive glycine pathway and the highly efficient enzymes that are unique to this pathway. These results demonstrate how process design strategies can enhance the feasibility of carbon-fixation systems, highlighting the potential of sequence optimization for improving the assimilation rate.

We performed the optimization for the same conditions as before, but imposed a larger upper bound on metabolite concentrations (*x*^ub^ = 0.15 M) to ensure that both pathways are feasible in the single- and multi-compartment configurations, so that their CO_2_ assimilation rates are comparable (Fig. 4A). As in the previous case, the reductive glycine pathway furnishes several orders-of-magnitude higher assimilation rates and its performance is less sensitive to enzyme saturation efficiencies than the Wood–Ljungdahl pathway in both reactor configurations. Moreover, for both pathways, optimizing the Michaelis constants is a more effective strategy to improve the assimilation rate in the multi-compartment configuration than it is in the single-compartment configuration.

**Figure 4:**
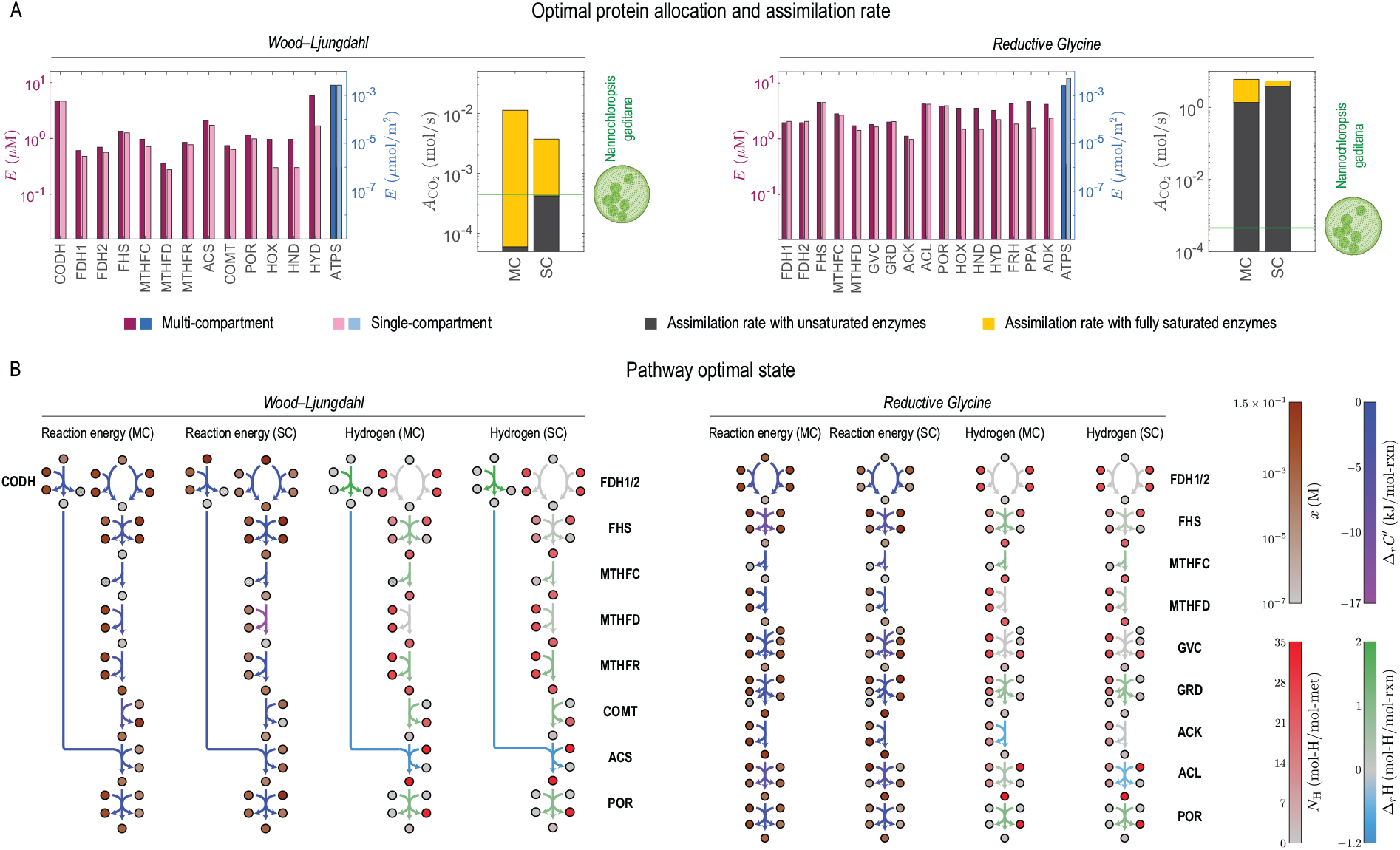
Optimal performance of C_1_-fixation systems in Fig. 3A. (A) Optimal protein allocation and CO_2_ assimilation rate, where pH_*c*_ = 7.5 for the single- and pH_*c*_ = 6 for the multi-compartment configuration (other parameters are reported in Table S1). Note that because no steady state can be achieved for the Wood–Ljungdahl pathway in the single-compartment configuration within physically relevant concentration ranges, a large upper bound on metabolite concentrations (*x*^ub^ = 0.15 M) is chosen for both pathways and configurations to optimize the carbon-fixation systems, so that the respective CO_2_ assimilation rates are comparable. Black bars indicate the assimilation rate if enzyme saturation efficiencies are accounted for, and yellow bars indicate how much the assimilation rate could be enhanced if all enzymes were fully saturated. (B) Optimal state of carbon-fixation pathways corresponding to the optimal design solutions in (A) for the single- and multi-compartment configurations. Pathway maps show the metabolite concentration *x*, transformed Gibbs free energy of reaction 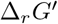, metabolite hydrogen content *N*_H_, and reaction hydrogen-ion consumption Δ_*r*_H.

Overall, optimal metabolite concentrations in the carbon compartment are such that the Gibbs free energy of reactions are as uniformly distributed along the carbon-fixation pathway as possible (Fig. 4B). Furthermore, pH-based compartmentalization is an effective design strategy for carbon-fixation systems, especially those constrained by strong thermodynamic bottlenecks, such as CODH in the Wood–Ljungdahl pathway. However, this strategy becomes less effective if there are no dominant thermodynamic bottlenecks, and the carbon-fixation pathway has thermodynamic anomalies (*i.e*., reactions that are less favorable in acidic environments), such as ACK and ACL in the reductive glycine pathway (Fig. 4B).

### Sensitivity to process and pathway parameters

We performed a sensitivity analysis to identify which parameter optimization strategies are the most impactful. From process parameters, we determined the sensitivity of the CO_2_ assimilation rate to the recycle ratios and found that increasing the recycle ratios in the range 0.1–0.5 leads to ∼1–10% improvement in the assimilation rate depending on the carbon-fixation pathway and reactor configuration (Fig. 5A). Granted, this strategy is more effective for the reductive glycine pathway since the baseline assimilation rate is much higher than for the Wood–Ljungdahl pathway.

**Figure 5:**
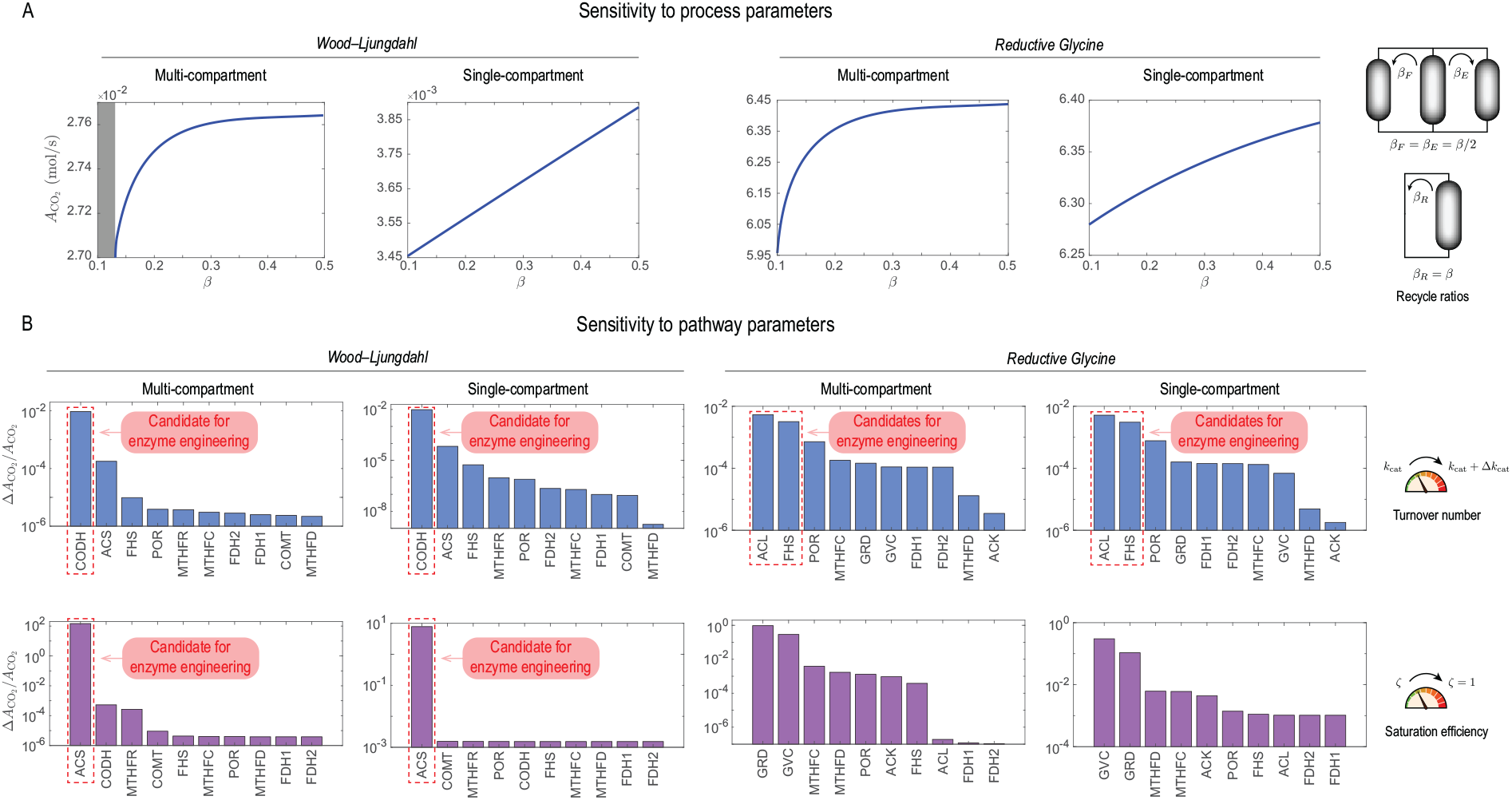
Sensitivity to pathway and process parameters. Sensitivity of optimal designs to (A) recycle ratios and (B) kinetic parameters. Shaded area in (A) indicates an infeasible range of the recycle ratios *β*_*F*_ and *β*_*E*_ for the Wood–Ljungdahl pathway in the multi-compartment configuration. Sensitivity to kinetic parameters is examined by perturbing the turnover number *k*_cat_ and saturation efficiency *ζ* of individual enzymes in each pathway and reactor configuration. The turnover number is increased by 1%, and the saturation efficiency is increased from an unsaturated level (*ζ <* 1) to fully saturated level (*ζ* = 1) of the respective enzyme.

From pathway parameters, we ascertained the sensitivity of the CO_2_ assimilation rate to the turnover number and enzyme saturation efficiency. Accordingly, improving the turnover number of CODH and ACS in the Wood–Ljungdahl along with ACL and FHS in the reductive glycine pathway has the largest impact on the assimilation rate for both reactor configurations (Fig. 5B). Enhancing the enzyme saturation efficiencies by tuning the Michaelis constants can also increase the assimilation rate. In this regard, ACS is the most effective target for enzyme engineering in the Wood–Ljungdahl along with CRD and GVC in the reductive glycine pathway for both reactor configurations (Fig. 5B).

### Physical limits of carbon fixation

As previously stated, C_1_ fixation is an energy-demanding process with slow kinetics. It is a reductive process predominantly fueled by reducing compounds. These general characteristics largely determine the design limits of C_1_-based manufacturing systems. Understanding these limits requires parametric studies of carbon-fixation systems. Since proton gradient and hydrogen are the energy sources that drive CO_2_ fixation in this work, we evaluated the CO_2_-assimilation capacity of the Wood–Ljungdahl and reductive glycine pathways as a function of proton and hydrogen concentrations.

When all other design parameters are fixed, the hydrogen concentration in the carbon compartment 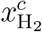 determines the rate and thermodynamic feasibility of redox regeneration reactions in both reactor configurations (see “Design optimization”). Similarly, the proton concentration in the carbon compartment, measured by pH_*c*_, controls the rate and thermodynamic feasibility of carboxylation reactions. Consequently, a trade-off between the thermodynamic driving forces of redox regeneration and carboxylation reactions arises at the pathway level, interacting with another between reactor sizes and stream flow rates at the process level. To globally capture these interacting trade-offs, we performed a parametric feasibility analysis of all the carbon-fixation systems considered in this study in the 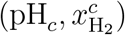 space.

The feasibility region for the Wood–Ljungdahl pathway in the single-compartment configuration is bounded from below in pH_*c*_ and 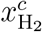 (Fig. 6A). The lower bound in pH_*c*_ corresponds to states where carboxylation (CODH) and ferredoxin regeneration (HYD) cannot both be thermodynamically feasible, while the lower bound in 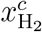 represents the minimum hydrogen concentration needed to drive the redox regeneration reactions at the same rate as the reductants are consumed by the carbon-fixation pathway. In contrast, the feasibility region in the multi-compartment configuration is bounded from above in pH_*c*_ and is unrestricted in 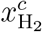 in the range we examined (Fig. 6B), suggesting that the hydrogen concentration is not a design-limiting factor.

**Figure 6:**
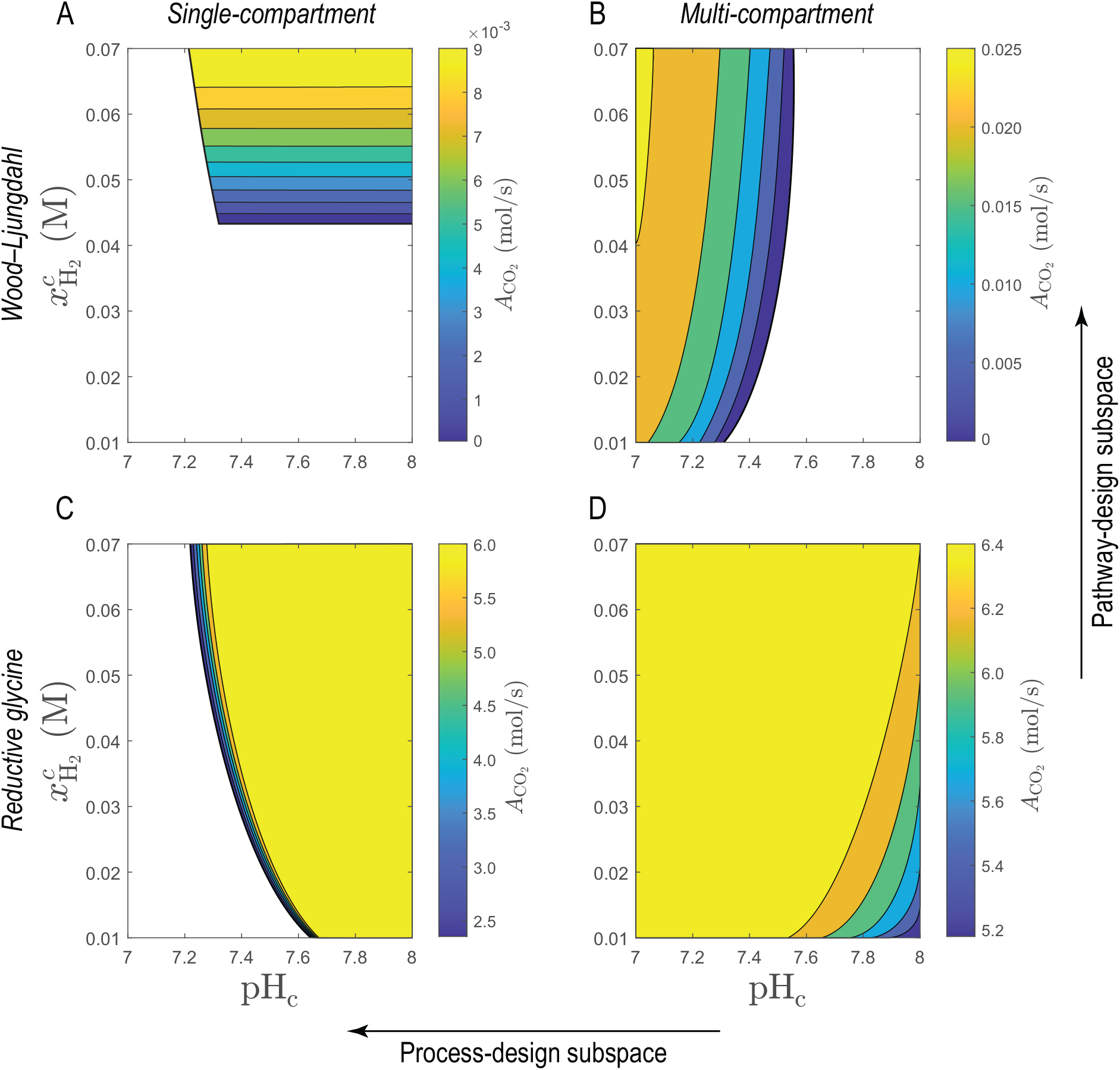
Feasibility of C_1_-fixation systems. Phase-portrait plots, showing the optimal CO_2_ assimilation rates for the Wood–Ljungdahl pathway in the (A) single-compartment and (B) multi-compartment configurations and for the reductive glycine pathway in the (C) single-compartment and (D) multi-compartment configurations. Black lines are the feasibility-region boundaries in the 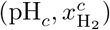-space for each case, where pH_*c*_ indicates the hydrogen-ion and 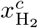 the hydrogen concentration in the carbon compartment. A large upper bound on metabolite concentrations (*x*^ub^ = 0.15 M) is imposed for both pathways and configurations to optimize the carbon-fixation systems, so that the respective CO_2_ assimilation rates are comparable (other parameters are reported in Table S1). All the enzymes are assumed to be fully saturated.

The feasibility region for the reductive glycine pathway in the single-compartment configuration is bounded from below in pH_*c*_, but is unrestricted in 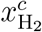 in the range we studied (Fig. 6C). As with the Wood–Ljungdahl pathway, carboxylation (POR) and ferredoxin regeneration (HYD) cannot both be thermodynamically feasible along this lower bound. However, the feasibility region in the multi-compartment configuration is not bounded in pH_*c*_ or 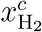 in the range we examined (Fig. 6D). Therefore, this combination of pathway and reactor configuration offers the greatest degrees of freedom for design optimization compared to other cases.

A few remarks concerning the design of the carbon-fixation systems discussed above are warranted. First, the feasibility region can be extended in both process and pathway design subspaces, highlighting the importance of coupling pathway and process design optimization. Second, from the four cases examined, the Wood–Ljungdahl pathway in the single-compartment configuration cannot operate in practice because the hydrogen concentrations required under reasonable physiological conditions (*x <* 0.05 M) to maintain a steady state is significantly higher than the hydrogen solubility. Last, the CO_2_ assimilation rate, reactor sizes, and stream flow rates can vary significantly across the feasibility region for the carbon-fixation systems studied. If economic objectives are to be optimized, what parameter sets from the feasibility region are optimal mostly depends on the operating and fixed costs associated with stream flow rates and reactor sizes.

## Discussion

Advancing towards a circular carbon economy requires innovative manufacturing strategies that efficiently utilize C_1_ carbon sources and are economically viable. However, designing C_1_-based manufacturing systems still remains a great challenge given that carbon fixation is a kinetically and energetically restricted process. Here, we developed a multi-scale, integrated systems approach to design C_1_-based manufacturing systems, coupling the optimization of pathway and process variables. We applied this approach in a biological, cell-free setting, examining the CO_2_-fixation capacity of the Wood–Ljungdahl and reductive glycine pathways in single- and multi-compartment reactor configurations. We showed that implementing the reductive glycine pathway in both reactor configurations results in several orders-of-magnitude higher CO_2_ assimilation rates than photosynthetic organisms, demonstrating the potential of the optimization framework we developed in identifying nontrivial manufacturing solutions.

The Wood–Ljungdahl and reductive glycine pathways, while vary similar, exhibit subtle, yet significant, differences in their stoichiometric and dissipative structures. These differences determine to what extent each pathway is restricted by thermodynamic constraints. Through parametric studies, we showed how these small differences could significantly impact the size of feasible design spaces for manufacturing systems in which these pathways are implemented. Overall, the reductive glycine pathway supports higher CO_2_ assimilation rates than the Wood–Ljungdahl pathway due to its comparatively uniform formation-energy landscape and its highly efficient enzymes, such as the glycine cleavage system [34–36]. The Wood–Ljungdahl pathway is an ancient reaction network [28] involved in energy conservation and biomass production [7]. In contrast, the reductive glycine pathway has likely co-evolved with lipid biosynthesis in modern organisms, coupling carbon and lipid metabolism [37, 38], leading to an evolutionary optimization of its unique enzymes. Extracting design principles from the structural characteristics of such evolutionary-tuned enzymes could guide the optimization of synthetic pathways for C_1_-based manufacturing in the future.

Cofactor regeneration is a major challenge for cell-free systems. The function of non-native C_1_-fixation pathways in cell factories relies on the redox and energy metabolism of the host. However, these pathways could disrupt native metabolism because they are energy-demanding, diminishing the growth rate of the host [25]. In contrast, cofactors must either be supplied or internally synthesized in cell-free systems. In this work, cofactors are regenerated by hydroganases and ATP synthase using hydrogen and proton gradient. Alternatively, these high-energy cofactors can be regenerated *via* the acid-aldehyde-ATP cycle using electricity as the energy source [39]. Integrating the acid-aldehyde-ATP cycle and carbon-fixation pathways in C_1_-based manufacturing systems could accelerate the electrification of industrial processes and support the renewable energy transition, making it a promising direction for future research.

The integrated systems approach developed in this study lays the groundwork for next generations of computer-aided design tools. Ideally, these generic tools can identify biotic, abiotic, or hybrid pathways for sustainable manufacturing to enable the production of value-added chemicals from diverse C_1_ feedstocks, offering economically viable solutions that support the transition towards a circular carbon economy.

## Methods

### Reactor systems

As stated in the main text, we decompose the design space of C_1_-based manufacturing systems into process and pathway design subspaces. We then examine two reactor configurations to assess the performance of carbon fixations systems: (i) A multi-compartment configurations with a redox, energy, and carbon compartments, and (ii) A single-compartment configuration with only a carbon compartment. These configurations may be regarded as representative process topologies from the process design subspace in which to search for optimal designs that maximize the energy efficiency and assimilation rate of carbon-fixation systems.

In the multi-compartment configuration, the enzymes of carbon-fixation pathways are placed in the carbon compartment, which operates under acidic conditions (pH_*c*_ *<* 7). Enzymes involved in redox regeneration are placed in the redox compartment under alkaline conditions (pH_*f*_ *>* 7). Similarly, enzymes involved in energy regeneration are placed in the energy compartment under neutral conditions (pH_*e*_ = 7). The volume of the redox, energy, and carbon compartments are denoted *V*_*f*_, *V*_*e*_, and *V*_*c*_, respectively. We treat all three compartments as continues stirred tank reactors (CSTR). The concentration of high-energy cofactors in the outlet stream *c* of the carbon compartment is always less than in the inlet *q*. Therefore, to replenish the high-energy cofactors consumed in the carbon compartment, a fraction of the outlet stream is directed to the redox and energy compartments. The recycle stream sent to the redox compartment for redox regeneration is denoted *F*, the flow rate of which is *ν*_*F*_ = *β*_*F*_ *ν*_*c*_, whereas the recycle stream sent to the energy compartment for ATP regeneration is denoted *E*, the flow rate of which is *ν*_*E*_ = *β*_*E*_*ν*_*c*_. Here, *ν*_*i*_ and *β*_*i*_ denote the flow rate and recylce ratio of process stream *i*. The concentrations of high-energy redox cofactors and ATP in the outlet stream of the redox and energy compartment, denoted *f* and *e*, are higher than in their respective inlet streams *F* and *E*. These energy rich streams are pumped back into the inlet stream of the carbon compartment to fuel carbon fixation. We denote the feed stream containing CO_2_ to the reactor system *Q* and the product stream containing pyruvate out of the reactor system *C*.

To regenerate the redox cofactors, aqueous hydrogen is pumped into the redox compartment at a rate 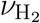 and concentration 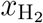. In the energy compartment, enzyme are partitioned into two groups: bulk and membrane enzymes. In this work, ATP synthase is the only membrane enzyme—others involved in energy metabolism are all bulk enzymes (see Fig. 2). Accordingly, ATP synthase units are distributed over a membrane with a surface area *A*_*e*_ and surface density *ρ*_*a*_. A highly acid fluid is pumped into the energy compartment through this membrane at a rate 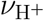 and concentration 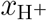 for ATP synthase units to regenerate ATP.

In the single-compartment configuration, enzymes involved in carbon metabolism, redox regeneration, and energy regeneration are all placed in the carbon compartment, which operates under near neutral conditions (pH_*c*_≈7.5) to ensure that both carboxylation and redox regeneration reactions are thermodynamically favorable. As with the multi-compartment configuration, the carbon compartment is regarded as a CSTR with the volume *V*_*c*_. Cofactors and unreacted metabolites in the outlet stream of the carbon compartment *c* are recycled back to the inlet stream *q* through a recycle stream *R*, the flow rate of which is *ν*_*R*_ = *β*_*R*_*ν*_*c*_. ATP is regenerated by ATP synthase units distributed over a membrane with a surface area *A*_*e*_ in the carbon compartment using a proton gradient as in the multi-compartment configuration. High-energy redox cofactors are also regenerated similarly by aqueous hydrogen, which is directly fed into the carbon compartment. As before, the feed and product streams of the reactor system are denoted *Q* and *C*, respectively.

### Design objectives

We seek two objectives to identify optimal strategies for designing C_1_-based manufacturing systems: maximize the (i) assimilation rate and (ii) energy efficiency. Multi-objective optimization problems are generally challenging to solve especially when their underlying constrains and objectives are highly nonlinear, as is the case in this work. To simplify this problem, we achieve the second objective by imposing the following energy-efficiency rules:

- Full redox recovery
- Full energy recovery
- Full CO_2_ recycling
- No excess redox regeneration
- No excess energy regeneration

The idea behind the formulation of these rules is two folds: One is to spend the minimum amount of energy needed to fix a given number of CO_2_ molecules, and the other is to minimize the circulation rates of recycle streams through the reactor systems examined in this study. The latter ensures that the residence time of reactants in the carbon compartment, where the thermodynamic and kinetic bottlenecks of the process reside, is maximized for a given volume *V*_*c*_.

The full redox and energy recovery rules imply that unreacted cofactors in the outlet stream of the carbon compartment are fully recycled, preventing loss through the product stream *C*. This recycling is crucial because cofactors are complex molecules and expensive to replenish. Full recycling maintains steady-state operation without additional supply, avoiding inefficient designs and unnecessary costs. These rules can be expresses as

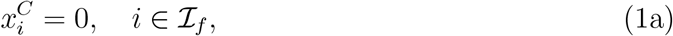

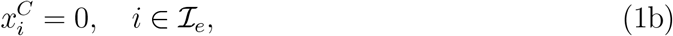

where 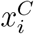 is the concentration of cofactor *i* in the product stream with ℐ_*f*_ and ℐ_*e*_ the index sets of redox and energy-transduction cofactors.

The full CO_2_ recycling rule ensures no CO_2_ loss through the product stream, maintaining a maximum concentration in the carbon compartment for a given amount of CO_2_ entering through the feed stream *Q*. Fixing the CO_2_ concentration and flow rate of the feed stream, this rule maximizes the flux through the carbon-fixation pathway. Consequently, it minimizes the volume of the carbon compartment needed to assimilate the total amount of CO_2_ pumped into the reactor system. The constraint representing this rule is

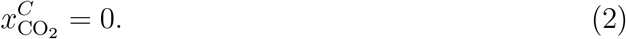

We impose additional constraints to balance the regeneration and consumption rates of redox and energy-transduction cofactors in the reactor system. These rules prevent cofactor loss through the product stream, improving energy efficiency and minimizing the required input of energy sources (H_2_ and H^+^) per CO_2_ fixed. Consequently, these rules reduce the circulation rate of materials through process streams by lowering the necessary input rates of H_2_ and H^+^. The constraints resulting from these rules can be categorized as either stoichiometric or global. We begin by describing the stoichiometric constraints first. The stoichiometric constraints ensure that cofactors are regenerated at the same rate as they are consumed by the carbon-fixation pathway. Unlike the previous rules, these constraints depend on the carbon-fixation pathway and reactor configuration.

For the Wood–Ljungdahl pathway in the multi-compartment configuration, we have

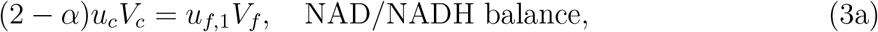

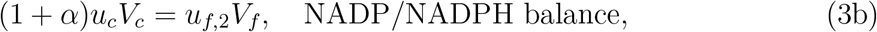

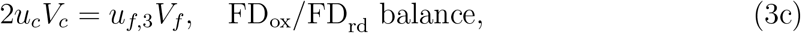

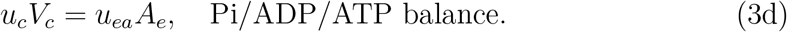

Here, *α* := *v*_FDH1_*/u*_*c*_ is a flux split ratio (see Fig. 3D) with *u*_*c*_, *u*_*f*,1_, *u*_*f*,2_, *u*_*f*,3_, and *u*_*ea*_ the canonical fluxes associated with the carbon-fixation pathway, NAD hydrogenase (HOX), NADP hydrogenase (HND), ferredoxin hydrogenase (HYD), and ATP synthase (ATPS), respectively (see Fig. 2). We will elaborate on canonical fluxes in section “C_1_-CAD framework”. Since carbon fixation, redox regeneration, and energy regeneration occur in the same reactor, the corresponding constraints in the single-compartment configuration are

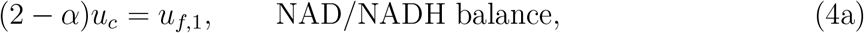

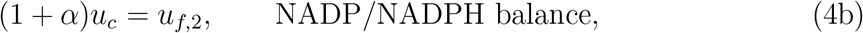

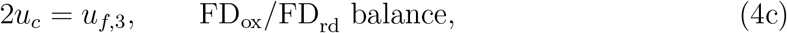

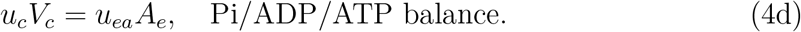

The redox- and energy-regeneration constraints for the reductive glycine pathway in the multi-compartment configuration are derived similarly

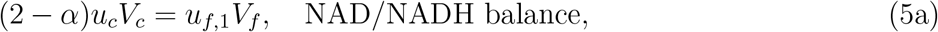

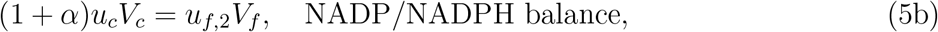

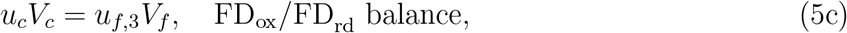

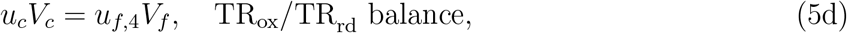

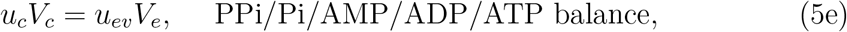

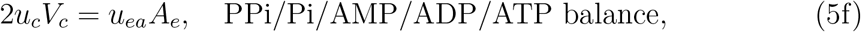

where *u*_*f*,4_ and *u*_*ev*_ are the canonical fluxes associated with thioredoxin hydrogenase (FRH) and the bulk enzymes of the energy compartment (PPA, ADK), respectively (see Fig. 2). The corresponding constraints for the single-compartment configuration are

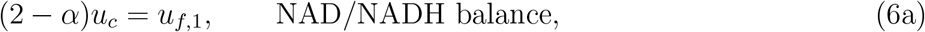

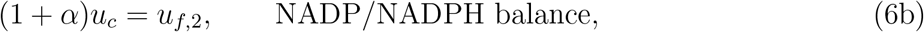

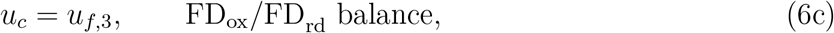

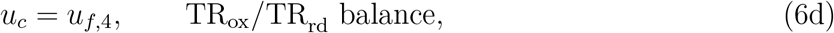

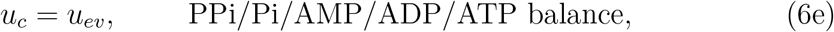

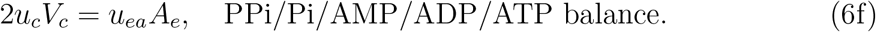

Next, we outline the global constraints. These constraints guarantee that the input rates of H_2_ and H^+^ match the energetic demands of the carbon-fixation pathway, as described below

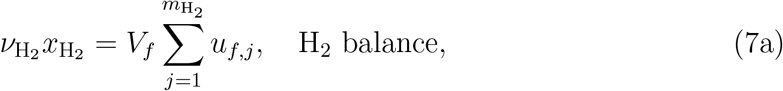

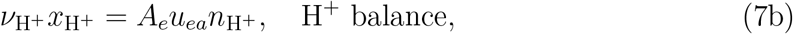

where 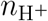 is the number of protons translocated per ATP molecule in ATP synthase, and 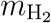 is the number of canonical fluxes in the redox compartment. We can derive a similar global balance for CO_2_ as follows

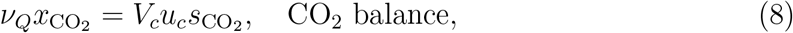

where *ν*_*Q*_ is the flow rate of the feed stream, 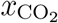 is the concentration of CO_2_ in the feed stream, and 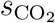 is the stoichiometric coefficient of CO_2_ in the carbon-fixation pathway.

### C_1_-CAD framework

We developed a Computer Aided Design framework to optimize C_1_-based manufacturing systems. This framework, termed C_1_-CAD, identifies optimal manufacturing solutions to produce industrial chemicals from C_1_ carbon sources by maximizing the objectives discussed in section “Design objectives” subject to the fundamental biophysical constraints of carbon fixation. The goal is to find optimal strategies for steady state and continuous manufacturing of industrial chemicals. The constraints imposed in this study are derived from first principles and include: (i) mass balance, (ii) kinetics, (iii) thermodynamics, (iv) ion binding, (v) Maxwell’s first law, and (vi) macromolecular crowding. In the following, we briefly outline these constraints, drawing on some of the techniques and formulations we previously developed [29, 32, 40–43].

We emphasize the importance of steady-state operation. Steady-state processes are generally safer, more predictable, easier to control, and cost less to maintain than alternative modes of manufacturing, providing greater consistency and product quality. Thus, we seek practical design strategies that ensure (i) steady-state manufacturing solutions can be achieved, and (ii) these solutions satisfy the foregoing fundamental biophysical constraints. In this study, solutions that satisfy these criteria are considered ‘feasible’; others are deemed impractical.

Mass-balance constraints apply both at the pathway and process level. At the pathway level, metabolic fluxes must be consistent with the stoichiometry of all the reactions in the entire carbon-fixation process at a steady state

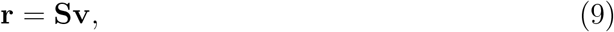

where **S** ∈ℤ^*n*×*m*^ is the stoichometry matrix, **v** ∈ ℝ^*m*^ is the flux vector, and **r** ∈ℝ^*n*^ the production-rate vector with *n* and *m* the number of metabolites and reactions in the entire process. Fluxes are decision variables of the C_1_-CAD framework to be determined by solving an optimization problem. To simplify this problem, we leverage the topology of carbon-fixation pathways to reduce the number of decision variables. Since both pathways we considered in this study are linear, we can characterize their flux sate by a single variable *u*_*c*_. We can also deduce from the stoichiometry of the cofactors that the fluxes associated with the bulk enzymes in energy regeneration can be characterized by a single variable *u*_*ev*_. However, the fluxes associated with redox regeneration can operate in parallel and, thus, can vary independently of each other. Therefore, the carbon-fixation fluxes **v**_*c*_ can be written

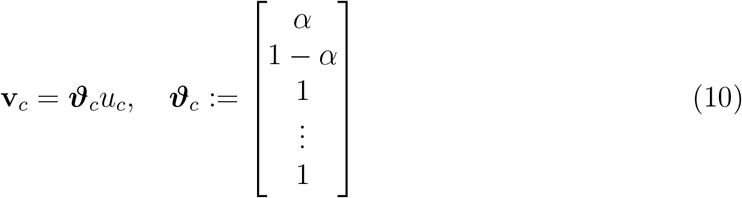

with the first two entries of ***ϑ***_*c*_ corresponding to FDH1 and FDH2, respectively. Accordingly, we refer to 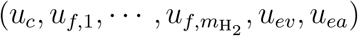 as canonical fluxes since they can vary independently without violating the stoichiometric constraints of all the reactions involved.

At the process level, we impose total and species mass-balance constraints at all process nodes. In the multi-compartment configuration, the total mass-balance constraints are

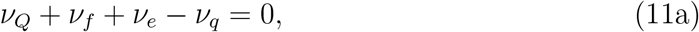

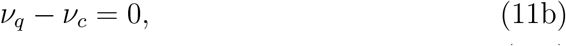

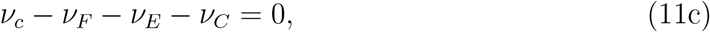

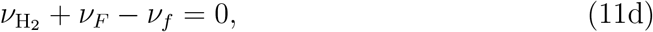

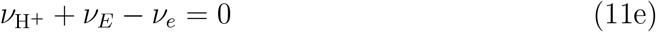

with the corresponding species mass-balance constraints for metabolite *i*

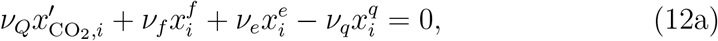

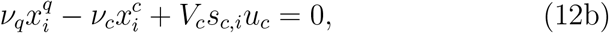

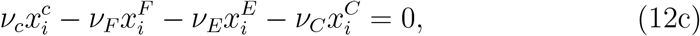

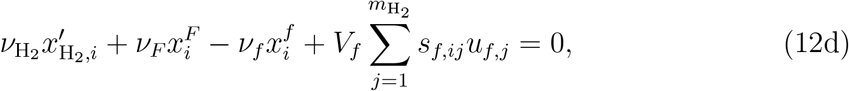

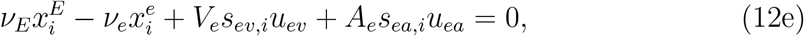

where

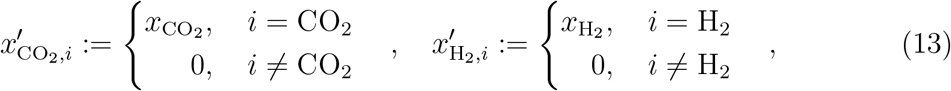

and 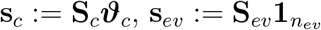. Here, **s**_*f,j*_ is the *j*th column of **S**_*f*_, **s**_*ea*_ corresponds to the ATPS column of **S**, 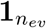 is an all-one column vector with *n*_*ev*_ entries, *n*_*ev*_ is the number of bulk enzymes in the energy compartment, and 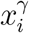 is the concentration of metabolite *I* in process stream *γ* with **S**_*c*_, **S**_*f*_, and **S**_*ev*_ the sub-matrices of **S** associated with carbon-fixation, redox regeneration, and bulk energy regeneration reactions, respectively. Note that metabolite concentrations in all process streams are generally low, not significantly affecting the density of water. Thus, we treated the density of water as a constant and express the total mass-balance constraints in terms of volume flow rates rather than mass flow rates in Eq. (11).

In the single-compartment configuration, the total mass-balance constraints are

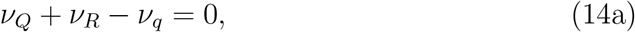

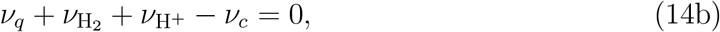

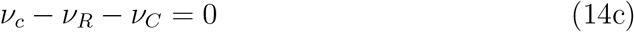

with the corresponding species mass-balance constraints for metabolite *i*

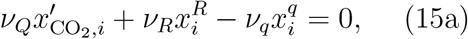

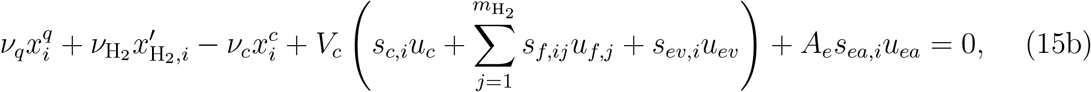

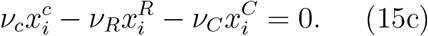

We impose additional constraints to specify how metabolites in the outlet stream of the carbon compartment are distributed between the recycle and product streams. In practical designs, a unit operation is needed to separate cofactros and unreacted intermediates of carbon-fixation pathway from the products. Here, to simplify the analysis, we specify the metabolite split ratios between the recycle and product streams without reference to any specific unit operation. In the multi-compartment configuration, the split ratios are given by

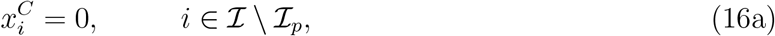

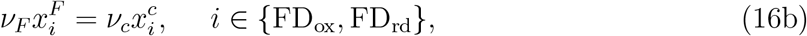

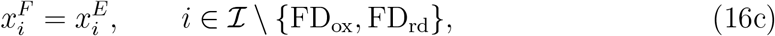

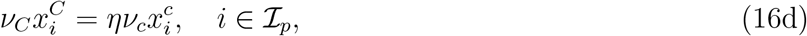

where ℐ is the index set of all the metabolites, ℐ_*p*_ is the index set of products (*i.e*., pyruvate), and *η* is the product recovery factor. Note that, in this work, we treat the FeS-clusters in ferredoxin-containing enzymes, such as FD_ox_ and FD_rd_, as metabolites. These clusters are water insoluble [44] and easier to separate than soluble cofactors, so we assume that they are completely separated from the outlet stream of the carbon compartment and sent only to the redox compartment, as described by Eq. (16b). However, for other metabolites, we assume no preferential separation, so they are uniformly distributed between the recycle streams *E* and *F*, as indicated by Eq. (16c). To enable a continuous regeneration of ferredoxins, the ferredoxin-containing enzymes circulate between the carbon and redox compartments, while other enzymes of carbon-fixation pathways are immobilized in the carbon compartment. For the products, the recovery factor indicates the amount of products that is recovered from the outlet stream of the carbon compartment and sent to the product stream. The unrecovered amount is then pumped back into the carbon compartment through the recycle streams. This split ratio is imposed by Eq. (16d). In the single-compartment configuration, the corresponding constraints are

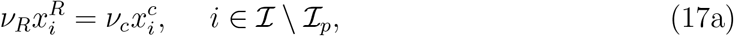

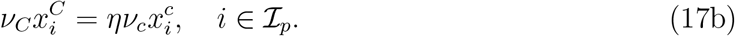

From the energy-efficiency and mass-balance constraints discussed above, we can derive a characteristic concentration, defined as

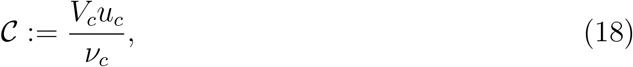

which, as will be discussed in section “Design optimization”, determines the right-hand side of equality and inequality constraints in the formulation of design-optimization problems for each reactor configuration. The characteristic concentration for the single- and multi-compartment configurations are given by (see Fig. 3C for definition of *s*_Pi_ and 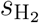)

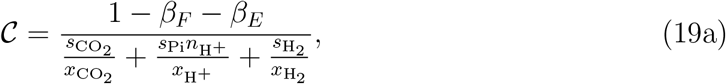

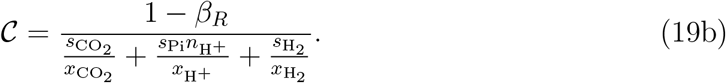

Applying the split-ratio constraints in Eqs. (16) and (17), we can express the product titer with respect to 𝒞 as

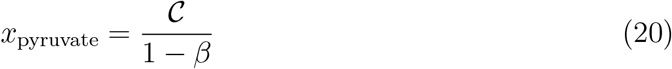

with *x*_pyruvate_ the concentration of pyruvate in the product stream and *β* the total recycle ratio, where *β* := *β*_*F*_ + *β*_*E*_ for the multi-compartment and *β* := *β*_*R*_ for the single compartment configuration.

Kinetic constraints are enzymatic rate laws that connect metabolic fluxes and metabolite concentrations. In this study, we apply the rate laws derived from the reversible Michaelis–Menten mechanism for all reactions except ATP synthase. This mechanism leads to the separable rate law [45]

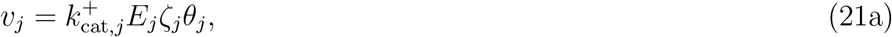

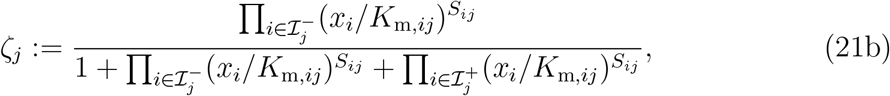

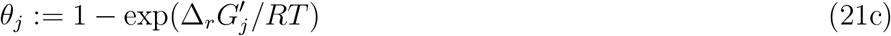

for reaction *j*. Here, *v*_*j*_ denotes the flux, *E*_*j*_ enzyme concentration, 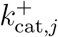 turnover number, *ζ*_*j*_ enzyme saturation efficiency, *θ*_*j*_ thermodynamic driving force, *K*_m,*ij*_ Michaelis constant for metabolite *i*, 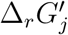 transformed Gibbs free energy of reaction,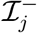 index set of substrates, and 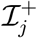 index set of products for this reaction with *R* and *T* the universal gas constant and temperature. For ATP synthase, the rate law is based on a rotary mechanism previously proposed [46], where the flux is expressed as a function of the proton gradient, membrane potential, and the concentrations of Pi, ADP, and ATP. The transformed Gibbs free energy of reaction is given by

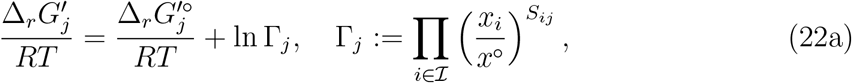

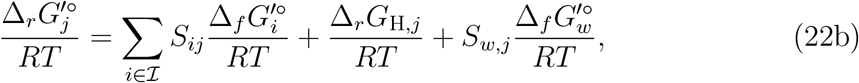

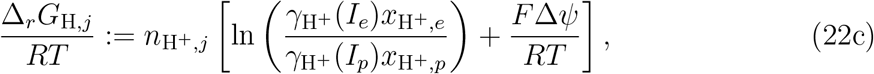

where Δ*ψ* is the membrane potential, *F* Faraday constant, *x*^°^ reference concentration, *S*_*w,j*_ stoichiometric coefficient of water, 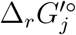 standard transformed Gibbs free energy of reaction, 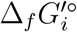 standard transformed Gibbs free energy of formation of metabolite *i*, 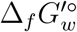 standard transformed Gibbs free energy of formation of water, *I* ionic strength, and 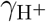 activity coefficient of hydrogen ion. Note that the subscripts *e* and *p* indicate that the corresponding parameters describe the conditions in the energy compartment and the acidic fluid pumped into the energy compartment, respectively. Here, the contribution of the membrane potential to 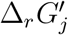 for membrane reactions in the energy compartment (ATPS) is accounted for by Eq. (22c) according to Maxwell’s first law [29]. Ion-binding constraints also contribute to 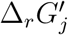, though implicitly through the formation energies in Eq. (22b) [32].

Whereas kinetic constraints concern stoichiometric mass balances, thermodynamic constraints determine reaction feasibility. The thermodynamic constraint are written as inequalities

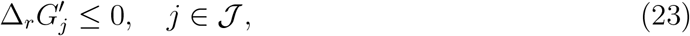

ensuring that metabolite concentrations are consistent with reactions proceeding in the reductive direction (forward directions shown in Fig. 2), where 𝒥 denotes the index set of all the reactions in carbon-fixation systems.

Macromolecular crowding constraints ensure that the amount of enzymes that can be packed in a unit volume of a reactor is bounded from above [47]. Typically, they are expressed with respect to the molar volume of enzymes. However, assuming that the densities of all the enzymes in the same compartment are approximately identical, we can express these constraints with respect to the molecular weights of enzymes. Accordingly, the macromolecular crowding constraints for the multi-compartment configuration are written

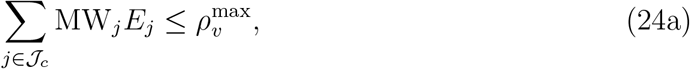

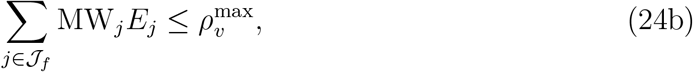

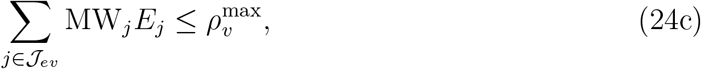

where 𝒥_*c*_ is the index set of carbon-fixation reactions, 𝒥_*f*_ the index set of redox-regeneration reactions, and 𝒥_*ev*_ the index set of bulk energy-regeneration reactions with MW_*j*_ the molecular weight of enzyme *j* and 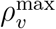 the maximum amount of enzymes (in grams) that can be packed in a unit volume (in liters). For the single-compartment configuration, we have

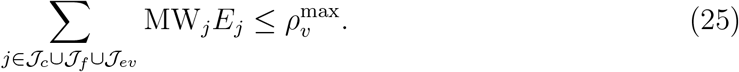

For membrane enzymes, a similar macromolecular crowding constraint is imposed

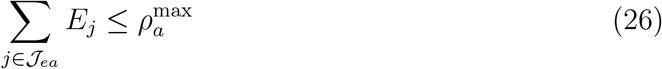

with 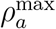 the maximum amount of enzymes (in moles) that can be packed in a unit membrane surface area (in m^2^). Note that, for membrane enzymes, the concentration *E*_*j*_ are measured in mol/m^2^, while for other enzymes, *E*_*j*_ are measured in M.

To impose the constraints outlined in this section, various types of biological data are required. Transformed Gibbs free energy of reactions were calculated using dissociation constants [48] and reference formation energies of species furnished by group-contribution methods [49], as discussed previously [32]. We generated manually curated datasets of experimentally measured kinetic and structural parameters for all the enzymes involved in the Wood–Ljungdahl and reductive glycine pathways along with those participating in energy and redox regeneration (Supplementary Data 1,2). All other parameters are reported in Table S1.

### Design optimization

We formulate an optimization problem that couples pathway and process design spaces to identify optimal manufacturing solutions for CO_2_ utilization. The objective is to maximize the CO_2_ assimilation rate 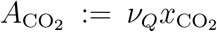 subject to the energy-efficiency and biophysical constraints discussed in previous sections. Accordingly, the objective we seek to maximize is *ν*_*C*_*x*_pyruvate_, noting that 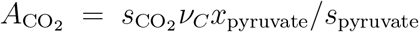. So far, we have partly simplified the analysis by incorporating energy-efficiency rules and canonical fluxes. However, solving the resulting optimization problem remains difficult, especially for the multi-compartment configuration. Therefore, we introduce additional techniques to further simplify the design optimization in this section. We first describe these techniques for the multi-compartment configuration in details, briefly discussing their counterparts for the single-compartment configuration at the end.

The first simplification technique focuses on optimizing the carbon compartment independently from other compartments, as it represents the main design bottleneck in the multi-compartment configuration. This approach involves expressing optimization constraints relative to metabolite concentrations in the carbon compartment outlet. After determining the optimal carbon compartment design, other decision variables are then adjusted to align with these optimized variables.

The second simplification technique incorporates the macromolecular crowding and kinetic constraints into the objective function. We can deduce from the energy-efficiency constraint Eq. (8) that maximizing *u*_*c*_ at a fixed *V*_*c*_ is equivalent to maximizing 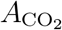. Given that the maximum *u*_*c*_ is achieved at a fixed *V*_*c*_ when the carbon compartment is fully packed with enzymes, it follows that the macromolecular constraint in Eq. (24a) must hold with equality. Accordingly, substituting the enzyme concentrations *E*_*j*_ from Eq. (21a) in Eq. (24a) and expressing **v**_*c*_ with respect to *u*_*c*_, as described by Eq. (10), we arrive at

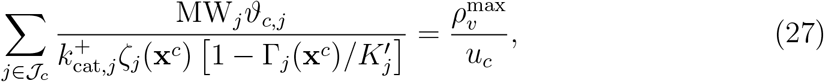

where

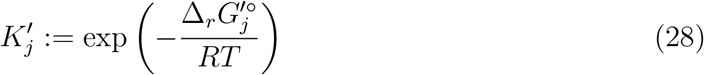

is the equilibrium constant of reaction *j*. We note that, to maximize *u*_*c*_ at a fixed *V*_*c*_, the left-hand-side expression in Eq. (27) must minimized with respect to the metabolite concentrations in the carbon compartment **x**^*c*^. Hence, the objective function to be minimized is

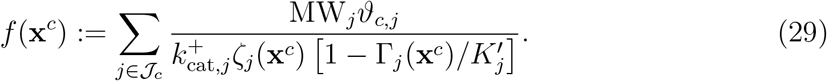

We minimize this objective for two cases: (i) The enzymes are all fully saturated, so that *ζ*_*j*_ = 1 for all *j*, and (ii) the enzymes are generally undersaturated (*ζ*_*j*_ *<* 1), and the saturation efficiencies are determined from the Michaelis constants and metabolite concentrations. The first case captures the physical limits of carbon fixation, while the second reflects the CO_2_-fixation capacity of the chosen pathway, given the structural properties and surface binding energies of the enzymes involved.

Having derived an objective function for the simplified design-optimization problem, we impose constraints on metabolite concentrations to restrict the design space. These constraints fall into three categories: (i) Cofactor ratios, (ii) upper and lower bounds, and (iii) thermodynamics. In the following, we briefly describe each category.

Cofactor-ratio constraints guarantee that the relative concentrations of high- and low-energy cofactors in the carbon compartment are such that they always provide a positive thermodynamic driving force for carbon fixation. These constraints are written as

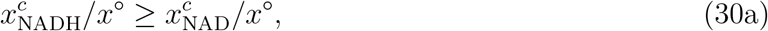

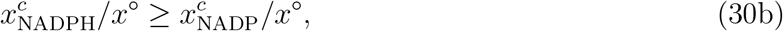

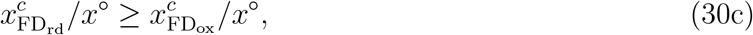

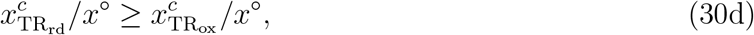

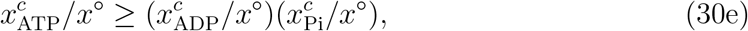

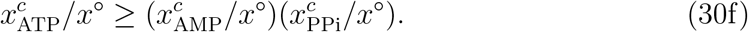

Note that Eqs. (30d) and (30f) apply only to the reductive glycine pathway, while the remaining equations apply to both pathways. We scale these constraints with the reference concentration *x*^°^ so that the left- and right-hand sides of the inequalities are dimensionally compatible.

Our goal in the general case is to find the optimal metabolite concentrations for all process streams that fall within physical range defined by upper and lower bounds, denoted **x**^lb^ and **x**^ub^, respectively. In the simplified design-optimization problem outlined above, these bounds can be directly imposed on metabolite concentrations in the outlet stream of the carbon compartment **x**^*c*^. However, to impose these bounds for other process streams, the respective metabolite concentrations must be expressed with respect to **x**^*c*^.These relationships are all linear and can be written as 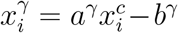 for metabolite *i* and process stream *γ* ∈ { *q, f, e, F, E* }. The constants *a*^*γ*^ and *b*^*γ*^ can be determined from the mass-balance constraints Eqs. (11) and (12), split-ratio constraints Eq. (16), and energy-efficiency constraints Eqs. (1)–(8). They depend on stoichiometric coefficients, recycle ratios, the concentration of boundary metabolites, and the characteristic concentration 𝒞. Applying the upper and lower bound constraints on metabolite concentrations to all process streams yields

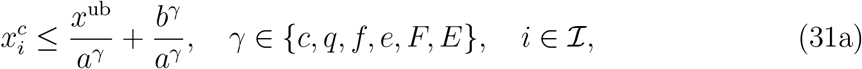

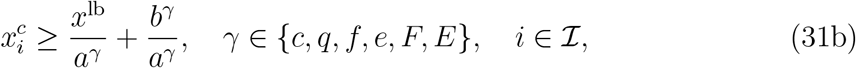

implying that

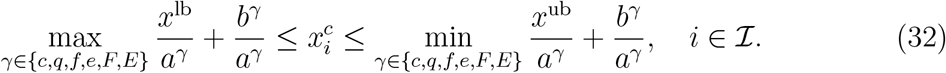

We impose these constraints using the same upper and lower bounds for all the metabolites in the carbon-fixation system.

The thermodynamic constraints Eq. (23) must hold in carbon, redox, and energy compartments to ensure that all reactions proceed in the forward direction. These constraints are evaluated with respect to the metabolite concentrations in the outlet stream of the respective compartment. For the carbon compartment, thermodynamic constraints can be expressed with respect to **x**^*c*^ explicitly as

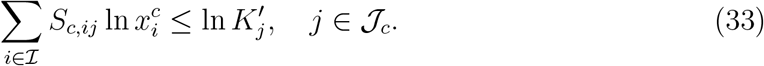

However, for the redox and energy compartments, thermodynamic constraints cannot be easily expressed with respect to **x**^*c*^ by logarithmically linear inequalities as in Eq. (33). To achieve a tractable formulation for the redox compartment, we express the thermo-dynamic constraints with respect to the concentrations of redox cofactors in the carbon compartment, assuming that the concentration of hydrogen in the carbon compartment 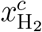 is a given parameter. We note that absolute value of 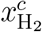 does not affect the mass-balance constraints of the carbon compartment since any unreacted amount of hydrogen from the redox compartment circulates through the reactor system. Therefore, Treating 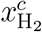 as a free parameter in the design-optimization problem is justified. Accordingly, the thermodynamic constraints of redox-regeneration reactions can be written as

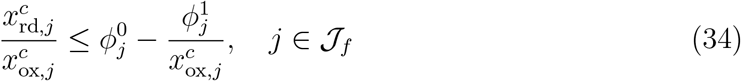

with 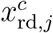 and 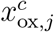 the concentrations of the high- and low-energy redox cofactors for redox-regeneration reaction *j* in the carbon compartment. Here, 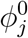 and 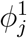 are constants that depend on 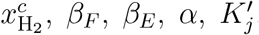, and C. Note that the ratio 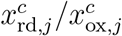 cannot be a negative number, implying that the right-hand-side of Eq. (34) must also be a non-negative number, from which we can derive the following concentration lower bounds

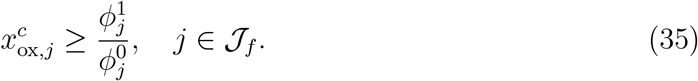

The lower bounds to be used for the concentrations of low-energy redox cofactors in the final design-optimization problem Eq. (39) is the maximum of the right-hand-side ratios in Eq. (35) and the corresponding lower bounds in Eq. (32). This lower bound is denoted 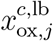.

The third simplification technique relates to the nonlinearity of Eq. (34). Because the thermodynamic constraints in this equation are logarithmically nonlinear in concentrations, we approximate the right-hand-side of Eq. (34) by piecewise log-linear functions. This is an appropriate approximation because the right-hand-sides are concave functions of 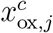 in the interval 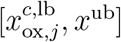. For this approximation, we introduce the auxiliary variables

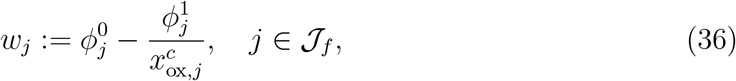

noting that ln *w*_*j*_ are also concave functions of ln 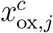. We then partition the interval 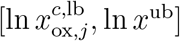 into *N*_*f*_ equally sized sub-intervals. In each of these sub-intervals, ln *w*_*j*_ is approximated as

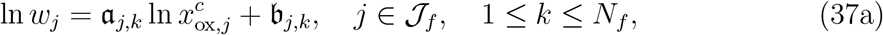

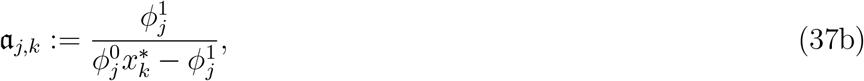

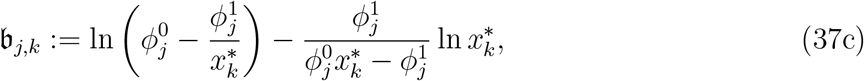

where 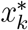 is a representative point from the *k*th sub-interval. Substituting these approximations in Eq. (34) yields

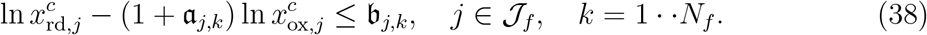

Note that the *N*_*f*_ inequalities in Eq. (38) for each reaction *j* are equivalent to the corresponding inequality in Eq. (34) as *N*_*f*_ → ∞.

A similar approximation technique can be applied to represent the thermodynamic constraints of the energy compartment with respect to **x**^*c*^. However, through numerical experiments, we found that the optimal concentrations furnished by the design-optimization problem generally satisfy these constraints without explicitly imposing them for the range of parameters we examined. Therefore, to simplify the analysis, we do not include these constraints in the final design-optimization problem.

We can now formulate a simplified optimization problem to identify optimal designs maximizing the assimilation rate and energy efficiency for the multi-compartment configuration. Representing all the constraints and objective with respect to **y**^*c*^ := ln **x**^*c*^, the simplified design-optimization problem is written

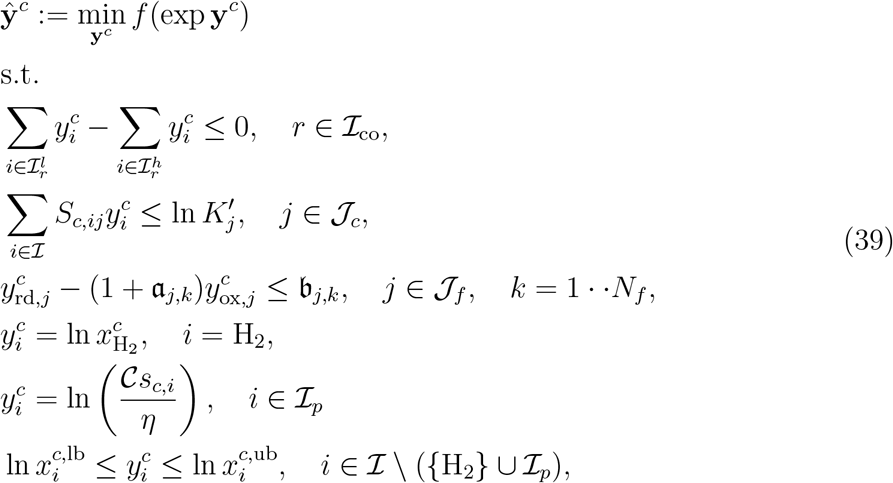

where 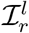 and 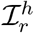 are the index sets of low- and high-energy cofacters associated with cofactor group *r* chosen from ℐ_co_. Note that, in Eq. (30), we refer to the cofactors contributing to each of the six cofactor-ratio constraints as a cofactor group; ℐ_co_ denotes the index set enumerating the six cofactor groups. Here, the objective function *f* (·) is given by (29) with 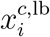and 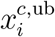 the lower and upper bounds on the concentration of metabolite *i* in the carbon compartment determined according to Eqs. (32) and (35). Once the optimal solution 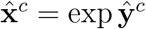 has been determined, the optimal canonical flux *û*_*c*_ can be ascertained from Eq. (27). From this optimal flux, the optimal enzyme concentrations **Ê**^*c*^ in the carbon compartment can then be determined using Eq. (21a).

Having identified the optimal operating conditions for the carbon compartment, the optimal design of the redox compartment is determined from the macromolecular-crowding constraint Eq. (24b) and specie mass-balance constraint Eq. (12d)

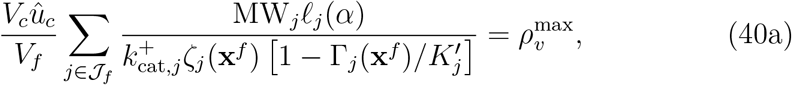

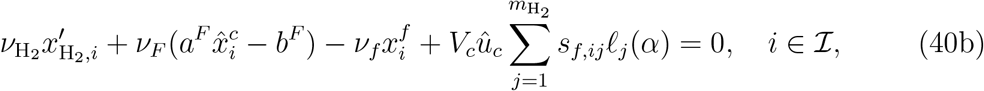

where the coefficients *ℓ*_*j*_(*α*) := *V*_*f*_ *u*_*f,j*_*/*(*V*_*c*_*u*_*c*_) are ascertained from the energy-efficiency rules Eqs. (3) and (5). Note that, in Eq. (40b), we have applied the linear relationship 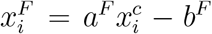, which we discussed earlier in this section. As with the carbon compartment, the optimal design is achieved when the reactor is fully packed with enzymes, so the macromolecular-crowding constraint Eq. (24b) must hold with equality. Eq. (40) provides a system of *n* + 1 equations, from which the optimal design variables 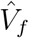 and 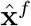 are to be determined.

We follow a similar procedure to identify the optimal design of the energy compartment. The rate law for the rotary mechanism of ATP synthase that we applied in this work [46] can be written as

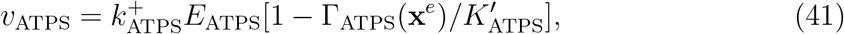

where 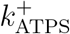 is a maximum turnover number determined from the parameters of the rotary mechanism. As with the redox and carbon compartments, the optimal design of the energy compartment is achieved when the reactor and membrane are fully packed with bulk and membrane enzymes, respectively. Accordingly, the macromolecular-crowding constraints Eqs. (24c) and (26) both must hold with equality. For the Wood–Ljungdahl pathway, the optimal design variables *Â*_*e*_ and 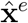 are the solution of

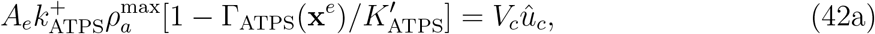

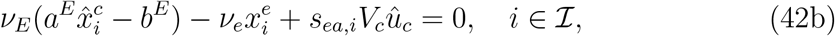

where we have applied the relationships 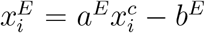 and 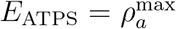 along with the energy-efficiency rule Eq. (3d). For the reductive glycine pathway, the optimal design variables are 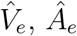, *Â*_*e*_, and 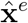, which are determined by solving

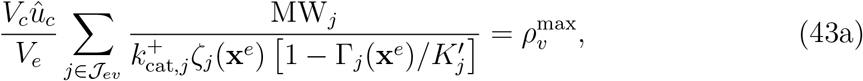

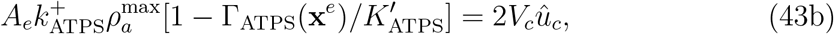

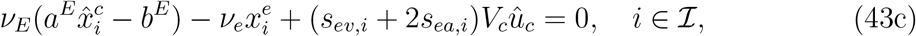

where we have applied the energy-efficiency rules Eqs. (5e) and (5f).

For the single compartment configuration, the cofactor-ratio constraints are the same as for the multi-compartment configuration. However, the upper and lower bounds are given by

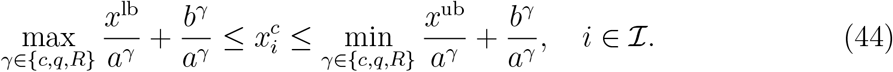

Moreover, the thermodynamic constraints for all reactions are expressed with respect to **x**^*c*^ as

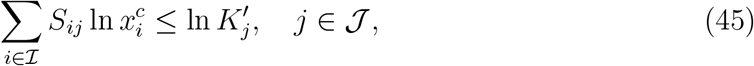

in which the equilibrium constants 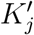 are evaluated at pH_*c*_. Accordingly, the simplified design-optimization problem is written as

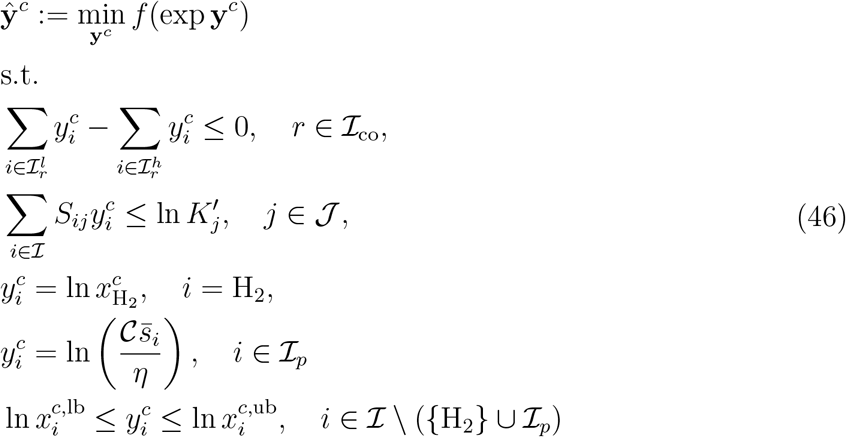

with the objective function

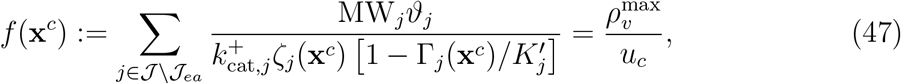

where

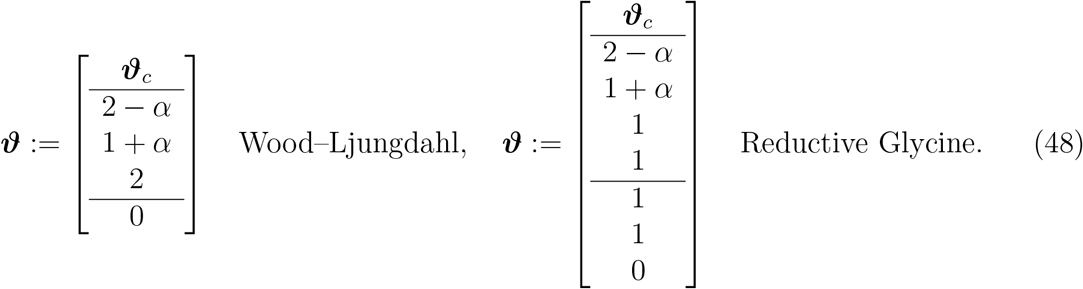

Here, 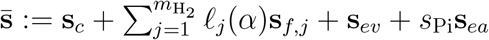 is a stoichiometry vector characterizing the production rates of metabolites in the single-compartment configuration (see Fig. 3 for definition of *s*_Pi_). Note that, in Eq. (48), the first, second, and third blocks correspond to the reactions involved in carbon metabolism, redox regeneration, and energy regeneration, respectively. The entries *ϑ*_*j*_ in the redox and energy regeneration blocks are determined from the stoichiometric relationships between the canonical fluxes in the energy-efficiency rules Eqs. (4) and (6). Once the optimal concentrations 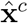 have been identified, the optimal flux *û*_*c*_ is ascertained from Eq. (47). Moreover, the optimal membrane surface area *Â*_*e*_ is determined from

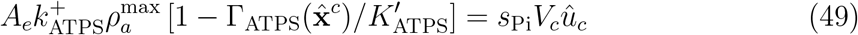

### Redox regeneration enzymes

Redox regeneration is a complex process in modern prokaryotes involving several enzymes. For example, NADH and NADPH are regenerated in *E. coli* by oxidizing high-energy carbohydrates through the Krebs cycle and pentose phosphate pathway [50], whereas reduced ferredoxins are regenerated by oxidoreductases and hydrogenase-mediated electron bifurcation using the energy from the oxidation of NADH and H_2_ [51, 52]. Incorporating these multi-step pathways for redox regeneration complicates the design of cell-free systems, rendering them costly and inefficient.

In this work, all redox cofactors are regenerated by hydrogenases. These ancient [53] and highly efficient [54] enzymes can directly regenerate the reduced cofactors by oxidizing H_2_, thus simplifying the design of cell-free systems. Although data on thioredoxin-specific hydrogenases are limited, recent studies have identified an frhAGB-encoded hydrogenase homologous to the F_420_-reducing hydrogenase in methanogens that can regenerate thioredoxins through direct oxidation of H_2_ [55, 56]. Accordingly, we estimated the CO_2_-fixation capacity of the reductive glycine pathway using the kinetic parameters of frhAGB (EC 1.12.98.1).

### Sensitivity to kinetic parameters

We examined the sensitivity of the CO_2_ assimilation rate to kinetic parameters to determine the best candidates for enzyme engineering in each carbon-fixation pathway. The desired improvement can be achieved through sequence optimization. Alternatively, natural enzymes with desirable kinetic properties can be identified *via* high-throughput screening. In most carbon-fixation pathways, carboxylation enzymes are usually the rate-limiting steps, and thus, they are the focus of enzyme screening studies. For example, metal-free formate dehydrogenases are extremely slow in the reductive direction [57], so they are undesirable candidates for the design of CO_2_-fixation systems. In contrast, metal containing formate dehydrogenases, such as those with molybdenum catalytic centers, are highly efficient, offering several orders-of-magnitude higher turnover numbers than their metal-free counterparts [58].

We examined the CO_2_-fixation capacity of the Wood–Ljungdahl and reductive glycine pathways using the kinetic parameters of metal-free and molybdenum-containing formate dehydrogenases. In this case study, the turnover number of the molybdenum-containing formate dehydrogenase was ∼10^5^ times that of the metal-free dehydrogenase. We found that, for the Wood–Ljungdahl pathway, this increase in the turnover number results in ∼ 10^1^ times higher CO_2_ assimilation rate (Fig. S1C), and, for the reductive glycine pathway, it results in ∼ 10^3^ times higher CO_2_ assimilation rate (Fig. S1D).

## Author Contributions

Conceptualization, A.A. and B.O.P.; Methodology, A.A.; Validation A.A.; Formal Analysis A.A.; Investigation, A.A.; Writing – Original Draft, A.A.; Writing – Review & Editing, A.A. and B.O.P.; Funding Acquisition, B.O.P.; Resources, B.O.P.; Supervision, B.O.P.

## Data Availability

All data generated or analyzed during this study are included in this published article and its Supplementary Information files.

## Code Availability

The source code used in this study is available online (https://github.com/akbari84/C1CAD.git).

## Acknowledgments

This work was funded by the Novo Nordisk Foundation (Grant Number NNF10CC1016517) and the National Institutes of Health (Grant Number GM057089). We thank Daniel Zielinski and Marc Abrams for their comments on the manuscript.

